# Speckle illumination microscopy enables slide-free and non-destructive pathology of human lung adenocarcinoma

**DOI:** 10.1101/2021.10.11.464016

**Authors:** Yan Zhang, Lei Kang, Claudia T. K. Lo, Terence T. W. Wong

## Abstract

Histopathology based on formalin-fixed and paraffin-embedded tissues remains the gold standard for surgical margin assessment (SMA). However, routine pathological practice is lengthy and laborious, failing to provide immediate feedback to surgeons and pathologists for intraoperative decision-making. In this report, we propose a cost-effective and easy-to-use histological imaging method with speckle illumination microscopy (i.e., HiLo). HiLo can achieve rapid and non-destructive imaging of large and fluorescently-labelled resection tissues at an acquisition speed of 5 cm^2^/min with 1.3-μm lateral resolution and 5.8-μm axial resolution, demonstrating a great potential as an intraoperative SMA tool that can be used by surgeons and pathologists to detect residual tumors at surgical margins. It is experimentally validated that HiLo enables rapid diagnosis of different subtypes of human lung adenocarcinoma and hepatocellular carcinoma, producing images with remarkably recognizable cellular features comparable to the gold-standard histology. This work will facilitate the clinical translations of HiLo microscopy to improve the current standard-of-care.

## Introduction

Histopathology has long term remained the gold standard for surgical margin assessment (SMA). However, routine pathological examination based on thin tissue slices (typically 5~10 μm) sectioned from formalin-fixed and paraffin-embedded (FFPE) tissues causes a significant delay in providing accurate diagnostic reports, thus failing to guide surgeons intraoperatively. Although frozen section can serve as an intraoperative diagnostic tool by cooling the tissue with the help of cryostat, it stills requires a turnaround time of 10 to 20 minutes which prolongs the operation duration. Besides, frozen sections are subjected to freezing artifacts when dealing with edematous and soft tissues, affecting slide interpretation and diagnostic accuracy^1^. It is reported that over 20% of cancer patients will undergo repeated surgeries^2^ due to the lack of a rapid and accurate tissue assessment tool that can provide immediate feedback to surgeons and pathologists for intraoperative decision-making.

A lot of efforts has been made to improve the routine clinical practice. Among them, optical imaging has demonstrated significant success in *in vivo* or *ex vivo* tissue imaging due to its rapid and non-invasive nature. Advanced microscopy techniques with optical sectioning capability enable to image a thin layer of a freshly excised tissue without the need for physically sectioning the specimen, greatly simplifying the procedures associated with slide preparation in conventional FFPE histology. The scanning-based depth-resolved imaging techniques, including reflectance and fluorescence confocal microscopy^3–5^, photoacoustic microscopy^6,7^, optical coherence tomography^8–10^, stimulated Raman scattering^11,12^, and nonlinear microscopy^13,14^, have demonstrated promising results in imaging human biopsied tissues. However, the requirement of sequential beam scanning limits the imaging throughput of these methods, posing a challenge to examine large resection specimens within a short diagnostic time frame. In contrast, wide-field imaging platforms which enable parallel pixel acquisition are advantageous for clinical applications due to their rapid imaging speed and reduced system complexity. Microscopy with ultraviolet surface excitation (MUSE)^15,16^, structured illumination microscopy (SIM)^17,18^, and open-top light sheet microscopy (OTLS)^19,20^ are all examples of this approach. MUSE relies on the limited penetration depth of ultraviolet light to achieve moderate optical sectioning, but it can exhibit some variations between different types of tissues^21^. OTLS is a versatile microscopy technique that enables rapid surface imaging and deep volumetric imaging of large specimens. However, oblique illumination will generate sever aberrations at the air-glass-water interface due to wavefront- and index-mismatching^22^, which have to be carefully corrected. Besides, axial resolution is inherently traded with filed-of-view (FOV) in Gaussian light sheet systems. In comparison, SIM enables digital rejection of out-of-focus background by leveraging the fact that only in-focus components can be modulated by the structured illumination. Although SIM is unable to obtain high-quality optical sections deep into the tissue^23^ since the grating contrast is rapidly deteriorated due to scattering, it is still a light-efficient imaging tool which has demonstrated clinical impact in diagnosis of prostate cancer^18,24^.

In this report, we propose a practical tissue scanner for intraoperative SMA with High-and-Low-frequency (HiLo) microscopy^25,26^. HiLo is a double-shot optical sectioning technique that relies on the acquisition of two images, one with speckle illumination and one with uniform illumination, eliminating the need for generating well-defined and controlled patterns as required by conventional SIM. In addition, HiLo is robust to aberrations and scattering in tissues as the statistics of fully-developed speckles are invariant^27^. We validate that HiLo can achieve rapid and non-destructive imaging of fresh mouse tissues at an acquisition speed of 5 cm^2^/min per fluorescence channel with a spatial resolution of 1.3 μm (lateral) and 5.8 μm (axial), which is competent to reveal subcellular diagnostic features and provide immediate on-site feedback. It is experimentally demonstrated that HiLo enables rapid diagnosis of different subtypes of human lung adenocarcinoma and hepatocellular carcinoma, producing images with remarkably recognizable cellular features comparable to the gold standard histology, showing its great potential as an assistive imaging platform for surgeons and pathologists to provide optimal adjuvant treatment intraoperatively. This work will facilitate the clinical translations of HiLo microscopy to improve the current clinical practice in pathological examination.

## Results

The HiLo system is configured in Epi-illumination mode (Fig. 1), allowing for convenient sample loading and accommodating tissues with arbitrary size and thickness. Two diode lasers that emit at 532 nm and 635 nm are utilized for simultaneous excitation of cytoplasmic and nuclear channel. TO-PRO3 is selected for nuclear staining since that it is a far-red DNA-selective dye which enables to be paired with other fluorophores for dual-channel fluorescence imaging without spectral crosstalk. The two coherent sources are combined by a dichroic beamsplitter and propagated through a variable beam expander, and subsequently illuminated on a rotatable diffuser to generate speckle and uniform illuminations required for HiLo imaging. The illumination patterns are focused onto the bottom surface of the specimen, which is translated in two scanning cycles, i.e., one with static diffuser for speckle illumination and the other with rapidly rotated diffuser for uniform illumination, for two-dimensional mosaic imaging. The excited fluorescence is collected by an inverted microscope, spectrally filtered and finally recorded by a monochromatic camera.

**Figure 1.**
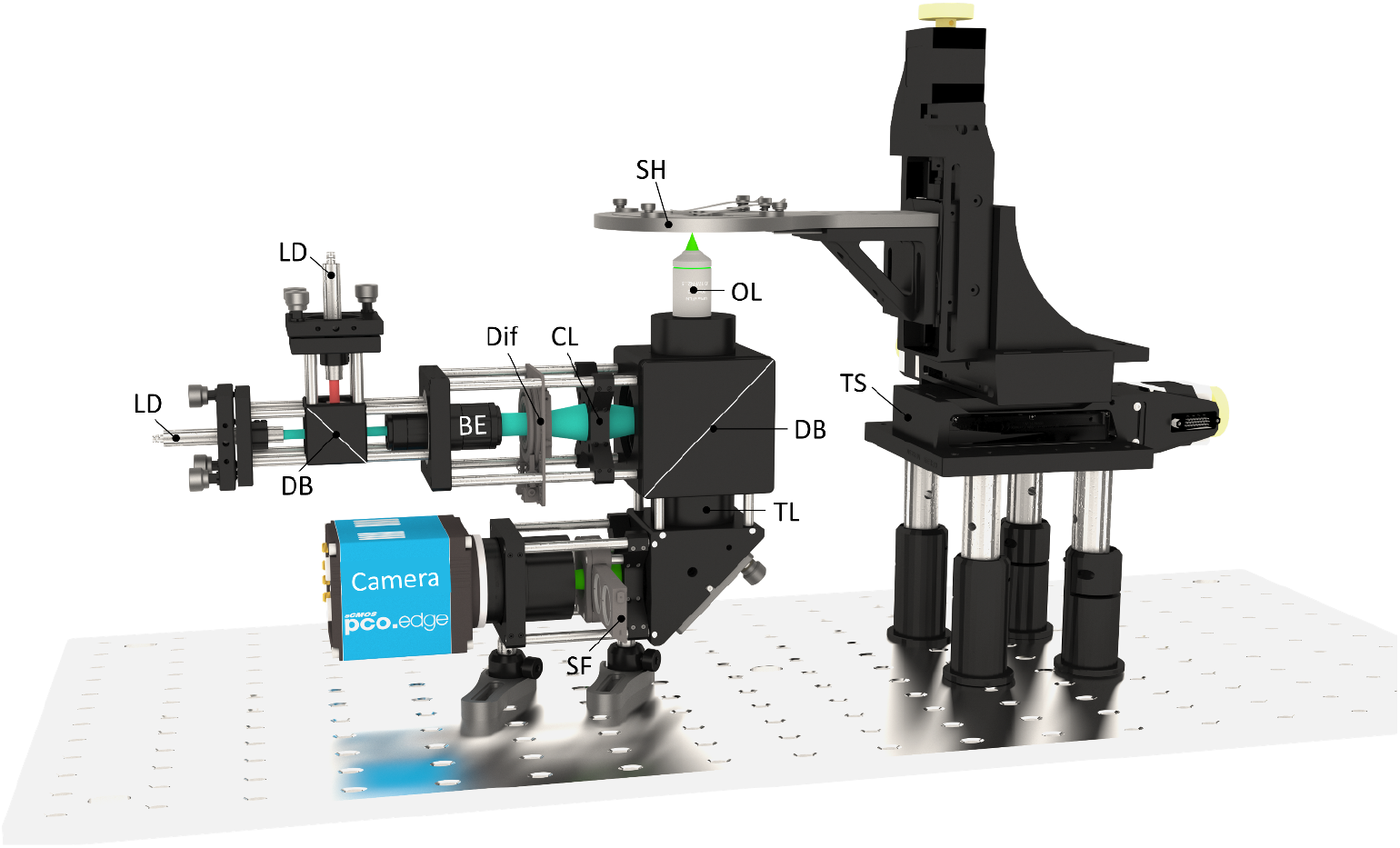
Schematic of the HiLo system for rapid histology-like imaging. Two laser diodes are combined by a dichroic beamsplitter and propagated through a variable beam expander, and are subsequently illuminated on a rotatable diffuser to generate uniform and speckle illuminations. The resulting illumination patterns are focused onto the bottom surface of the specimen through a condenser lens and an objective lens. The excited fluorescence from cytoplasmic and nuclear channel is successively collected by the same objective lens, refocused by a tube lens, spectrally filtered, and finally imaged by a monochromatic camera. The specimen is supported by a sample holder and raster-scanned by motorized stages. LD, laser diode; DB, dichroic beamsplitter; BE, beam expander; Dif, diffuser; CL, condenser lens; OL, objective lens; TL, tube lens; SF, spectral filter; SH, sample holder; TS, translational stage.

It has been reported that strong scattering in thick tissues poses a challenge for HiLo imaging by reducing the in-focus speckle contrast^28^, which serves as a weighting function to numerically filter the out-of-focus contributions of the specimen. This effect can be alleviated by illuminating with a coarse speckle pattern at the cost of a compromised optical sectioning strength. A mouse brain is utilized to determine the optimal speckle frequency for HiLo imaging of thick and scattering tissues. A formalin-fixed brain is sectioned at the surface by a vibratome to obtain a thin tissue slice (10-μm thickness) and a thick tissue block (~5-mm thickness), and both of them are stained with 10 μM TO-PRO3 for 1 min and rinsed in phosphate-buffered saline (PBS) for 30 s. The tissues are illuminated by speckle patterns with varying frequencies by changing the beam diameter illuminated on the diffuser, and subsequently imaged by an objective with a numerical aperture (NA) of 0.3. The speckle contrast *C_s_* is calculated as the ratio of standard deviation to mean of the speckle-illuminated images (*I_s_*). The quantitative results are plotted in Fig. 2. As expected, the speckle contrast will quickly degrade with the increase of speckle frequency (*f_s_*), and this effect is even worse in thick tissues which can cause an underestimation of the weighting function even though the specimen is in focus. Besides, it is proved that for a given sample thickness, *C_s_* is lower for high-NA objective since its narrow depth-of-field (DOF) will further exacerbate the out-of-focus fluorescence^28^. To make a good balance between sectioning capability and signal-to-noise ratio (SNR) of the HiLo images, a 10×/0.3NA objective lens is implemented, and the speckle frequency is set to 250 mm^-1^ in this study, which is corresponding to an averaged speckle size ~4 μm.

**Figure 2.**
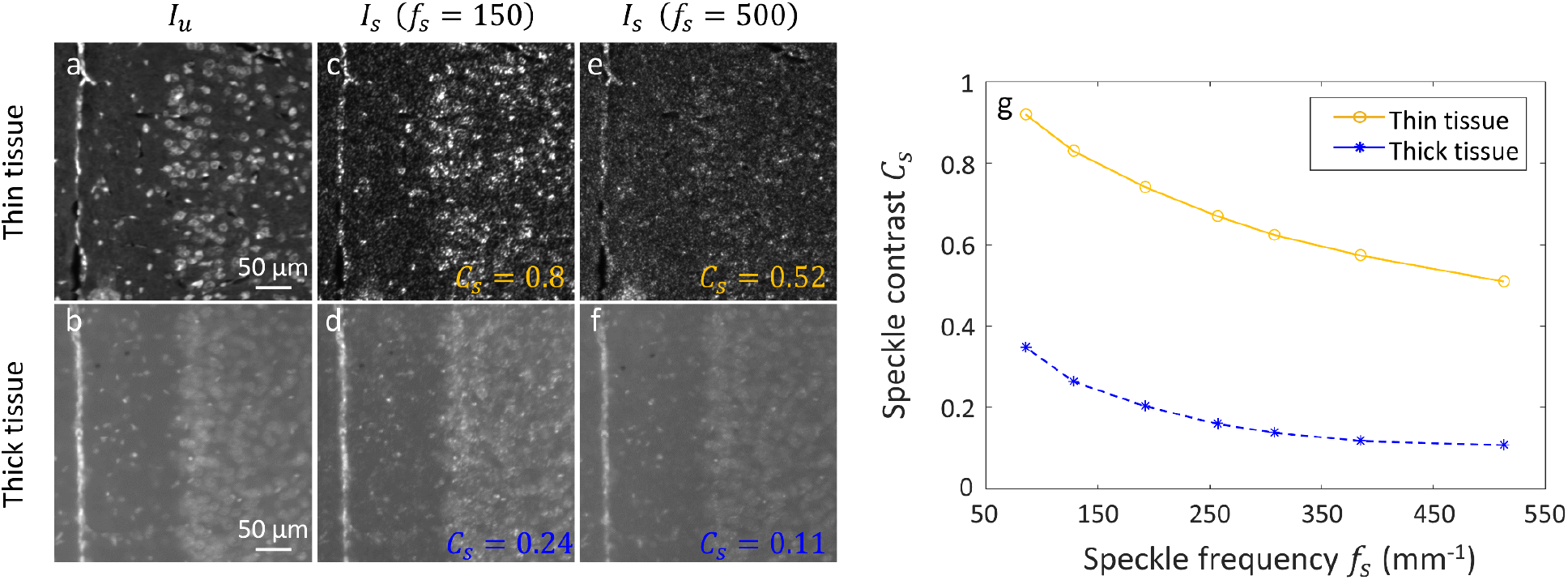
Experimental determination of the optimal speckle frequency for HiLo imaging of thick tissues. **a,b,** Uniformly-illuminated images of thin (~10-μm thickness) and thick (~5-mm thickness) mouse brain tissues, respectively. **c,d,** Speckle-illuminated images of thin and thick mouse brain tissues with a speckle frequency of 150 mm^-1^. **e,f,** Speckle-illuminated images of thin and thick mouse brain tissues with a speckle frequency of 500 mm^-1^. **g,** The relationship between speckle frequency and speckle contrast that can be observed.

Fluorescent beads with a diameter far below the resolution limit of a microscope is commonly used to experimentally determine the system’s axial resolution. However, this will collide with the filtering process in HiLo microscopy which achieves optical sectioning by evaluating the speckle contrast over a sampling window containing several imaged grains. Alternatively, we quantify HiLo’s axial resolution by imaging 10-μm-diameter fluorescent microspheres^29^, and the resulting axial resolution can be calculated as:

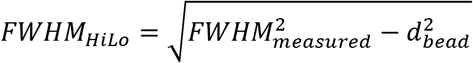

where *FWHM_measured_* is the full width at half maximum (FWHM) of the measured optical sectioning curve and *d_bead_* is the diameter of fluorescent microspheres, which is 10 μm according to the manufacturer. The specimen is axially scanned over a total range of 50 μm with a step of 0.5 μm. The HiLo images (Fig. 3b–e) and wide-field images (Fig. 3f–i) at different axial scanning depths are compared in Fig. 3. The curve of axial intensity distribution of a selected fluorescent microsphere (indicated by orange box in Fig. 3b–e) is shown in Fig. 3j, with a measured FWHM of 11.6 μm, corresponding to an axial resolution of 5.8 μm, which is competent to produce an optical section in lieu of a physical section in slide-based FFPE histology. In addition, different sectioning capability can be obtained from the same raw dataset by adjusting an optical section parameter σ (see methods) during numerical processing^26^ (Fig. 3k), which is the unique feature of HiLo microscopy. The image is sampled at 0.65 μm/pixel with 10× magnification, leading to a Nyquist-limited lateral resolution of 1.3 μm. Despite this, HiLo is still capable of revealing subcellular features which are essential for diagnosis.

**Figure 3.**
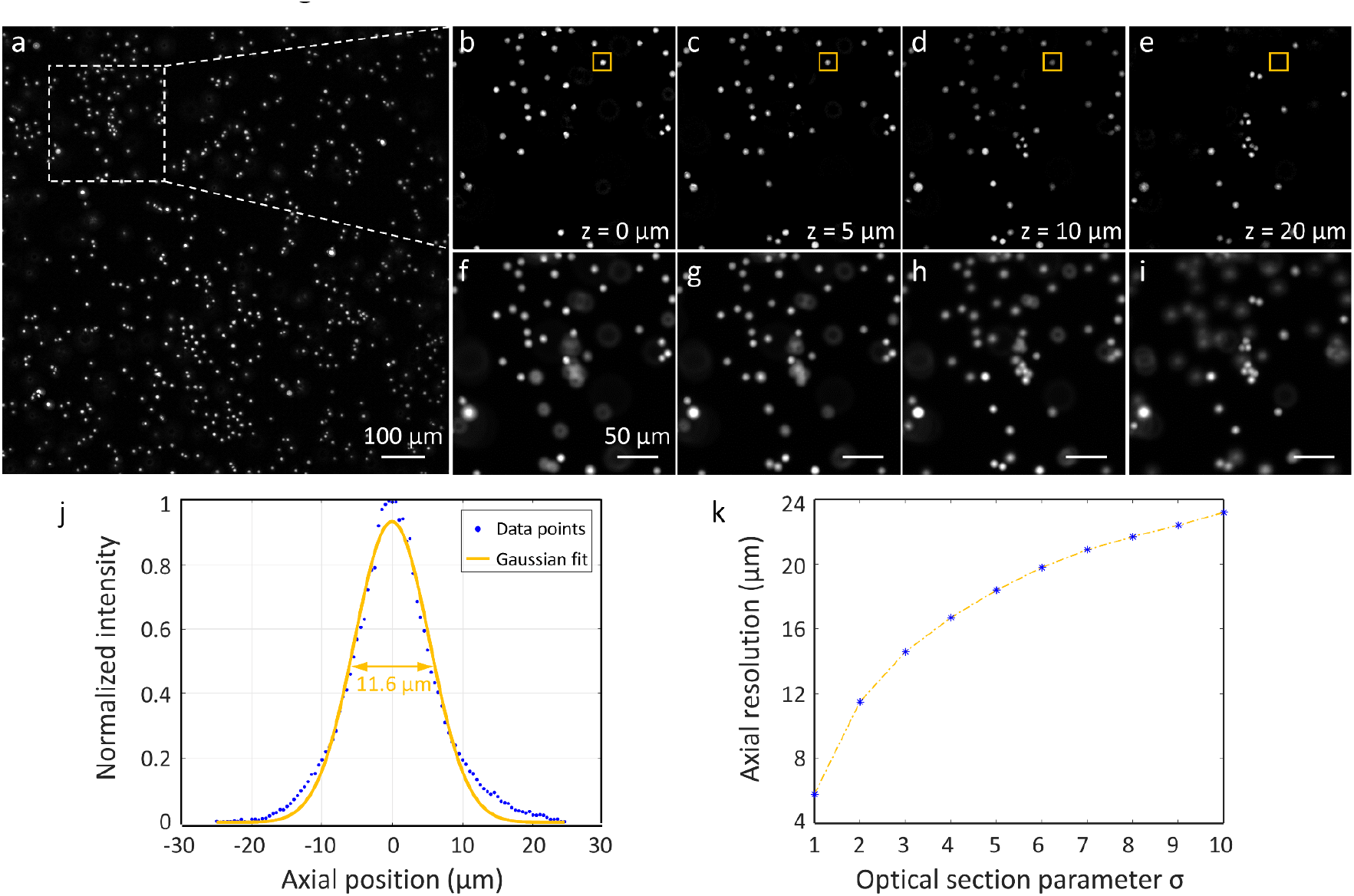
Experimental characterization of HiLo’s axial resolution. **a,** Maximum-intensity projection of the HiLo images of 10-μm-diameter fluorescent microspheres. **b–e,** Zoomed-in HiLo images of white dashed region in a at different axial depths. **f–i,** The corresponding uniformly illuminated wide-field images. **j,** Axial intensity distribution of a selected microsphere (indicated by orange box in b–e) with an axial step of 0.5 μm. **k,** The relationship between axial resolution and optical section parameter σ. The experiment was performed with a 10×/0.3NA objective at 532-nm excitation wavelength and 560-nm fluorescent emission wavelength.

Freshly excised mouse tissues, including heart (Fig. 4a–f), brain (Fig. 4g–l), kidney (Fig. 4m–r), and liver (Fig. 4s–x), are manually sectioned with a thickness of ~5 mm by a scalpel and labeled by TO-PRO3, and subsequently imaged by HiLo to validate the system’s performance initially. The top row of each color panel shows the uniformly-illuminated wide-field images whereas the bottom row represents the HiLo images, in which the densely packed cell nuclei can be clearly resolved with a significantly improved image contrast. The camera integration time for each raw fluorescence image is generally less than 100 ms in our experiments, corresponding to a minimum acquisition speed of 5 cm^2^/min per fluorescence channel.

**Figure 4.**
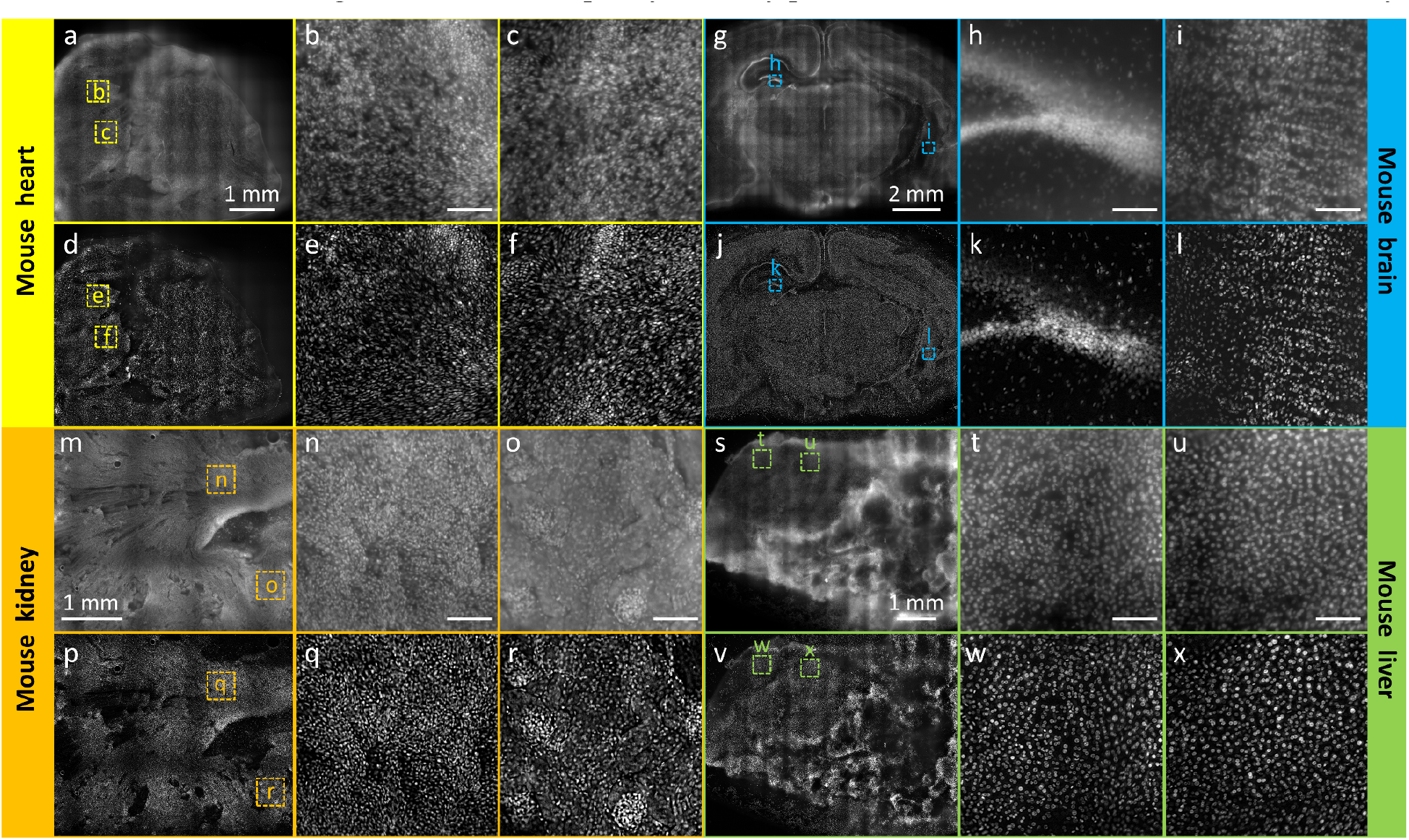
HiLo imaging of freshly excised mouse tissues. **a–f,** HiLo images of mouse heart. **g–l,** HiLo images of mouse brain. **m–r,** HiLo images of mouse kidney. **s–x,** HiLo images of mouse liver. The top row of each color panel are uniformly-illuminated wide-field images whereas the bottom row are optically sectioned HiLo images. Scale bar: 100 μm.

TO-PRO3 and eosin can serve as a fluorescent analog to hematoxylin and eosin (H&E) staining, enabling dual-channel fluorescence imaging which could be false-colored to mimic the H&E-stained histological slides^30^. To showcase, a fresh mouse heart (~5 mm thickness) is submerged in 2% eosin and 10 μM TO-PRO3 in PBS, and successively imaged by HiLo to produce dual-channel images, which are then false-colored by a previously reported method^31^ (Fig. S1). We and other groups found that eosin tends to leak out of the tissue during imaging since it is weakly bound to fresh and hydrated specimens^20,32^, generating unwanted background fluorescence which deteriorates the image contrast. For simplicity, we only preserve nuclear channel in this validation study.

To demonstrate the full potential of HiLo in the application of intraoperative SMA, formalin-fixed but unprocessed human lung tissues (Fig. 5) and human liver tissue (Fig. 6) are imaged. The specimens are fixed in formalin immediately after surgical excision to prevent tissue degradation. After HiLo imaging, the specimens are processed following the standard protocol (i.e., paraffin embedding, microtome sectioning, slide mounting and staining) to obtain the H&E-stained histological images for comparison. Note that H&E-stained thin tissue sections are not able to exactly replicate the surface imaged by HiLo since that tissues can be distorted during the preparation procedures.

**Figure 5.**
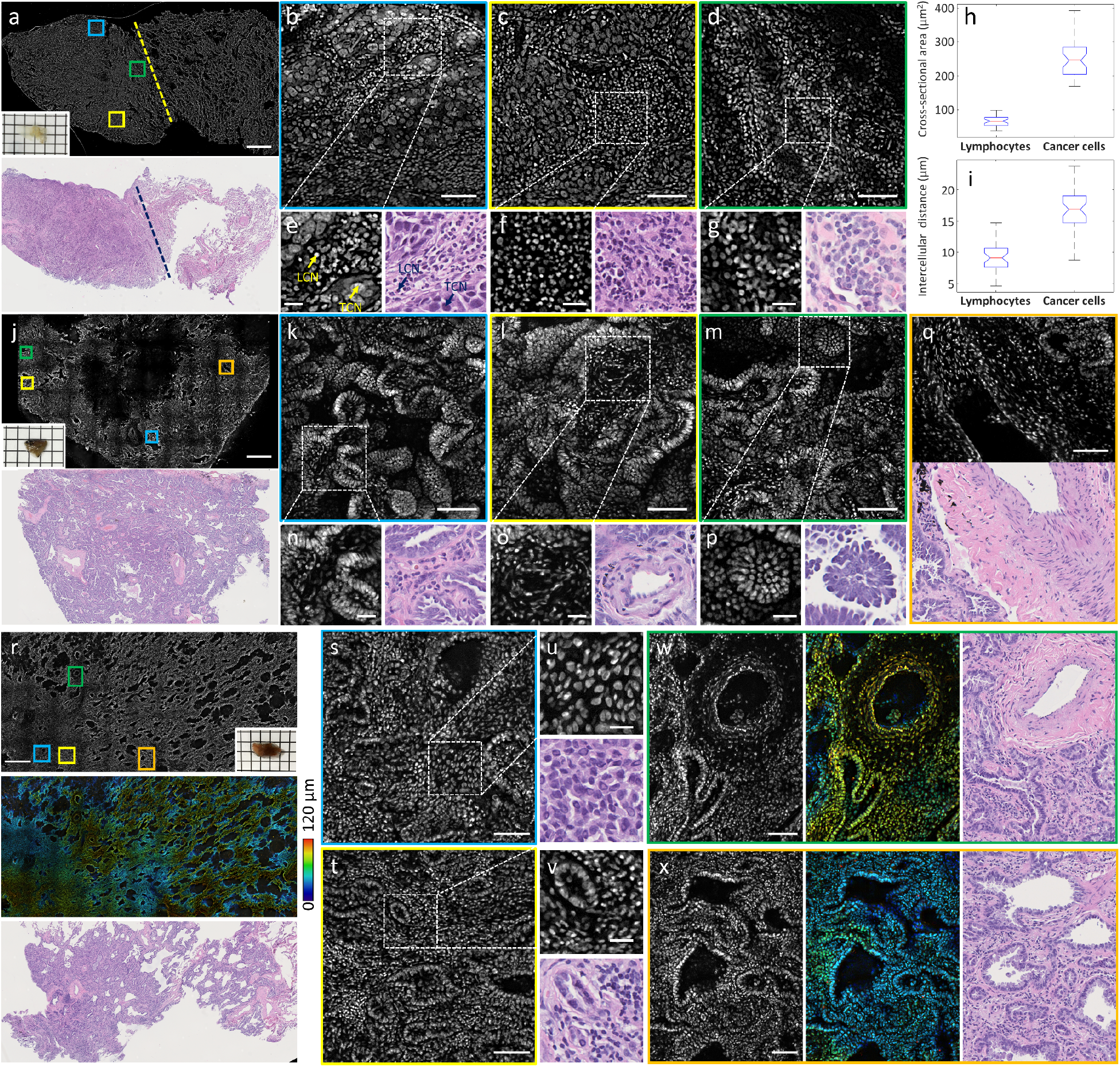
HiLo imaging of human lung tissues. **a**, HiLo (top) and H&E-stained image (bottom) of a fixed human lung specimen with large cell carcinoma, the dashed lines outline the boundary between normal (right) and tumour (left) regions. Inset at the bottom left of HiLo shows the photograph of the specimen. **b–d,** Zoomed-in HiLo images of blue, yellow, and green solid regions in a, respectively. **e–g,** Zoomed-in HiLo and H&E-stained images of white dashed regions in b–c, respectively. LCN, lymphocyte cell nucleus; TCN, tumor cell nucleus. **h,** Distribution of nuclear cross-sectional areas of lymphocytes and cancer cells derived from b (*N* = 50) with a median of 65 μm^2^ and 240 μm^2^, respectively. **i,** Distribution of intercellular distances of lymphocytes and cancer cells derived from b (*N* = 50), with a median of 9 μm and 16 μm, respectively. **j**, HiLo (top) and H&E-stained image (bottom) of a fixed human lung specimen with papillary-predominant adenocarcinoma, inset at the bottom left of HiLo shows the photograph of the specimen. **k–m,** Zoomed-in HiLo images of blue, yellow, and green solid regions in a, respectively. **n–p,** Zoomed-in HiLo and H&E-stained images of white dashed regions in k–m, respectively. **q,** Combined HiLo and H&E-stained mosaic image of orange solid region in a. **r**, HiLo (top), surface topology (middle), and H&E-stained image (bottom) of a fixed human lung specimen with acinar-predominant adenocarcinoma, inset at the bottom right of HiLo shows the photograph of the specimen. **s,t,** Zoomed-in HiLo images of blue and yellow solid regions in a, respectively. **u,v,** Zoomed-in HiLo and H&E-stained images of white dashed regions in s and t, respectively. **w,x,** Zoomed-in HiLo (left), surface topology (middle), and H&E-stained images (right) of green and orange solid regions in r, respectively. Scale bar, 1 mm (a,j,r), 30 μm (e–g,n–p,u,v), and 100 μm (the rest).

**Figure 6.**
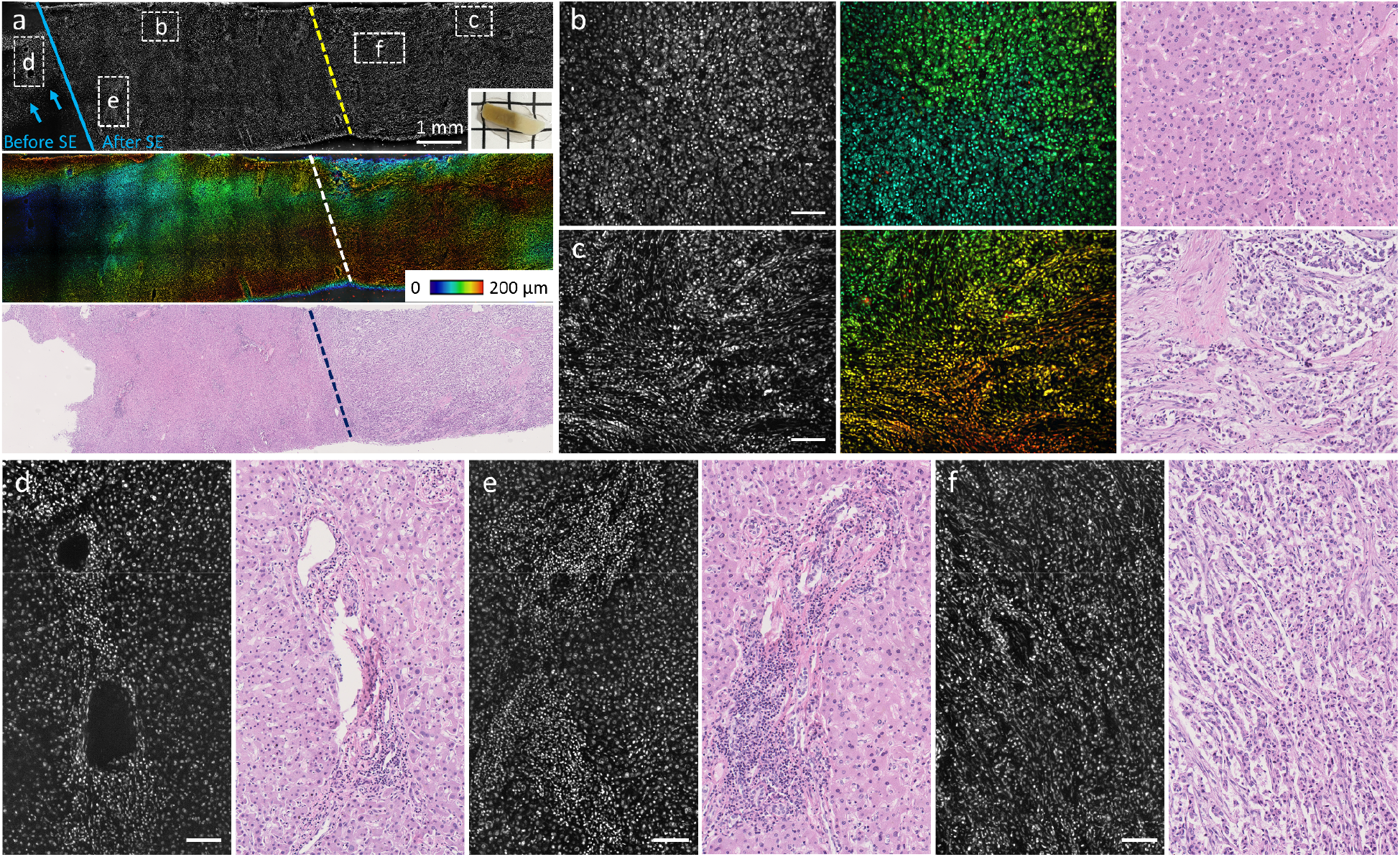
HiLo imaging of a human liver tissue. **a**, HiLo (top), surface topology (middle), and H&E-stained image (bottom) of a fixed human liver specimen with scirrhous-subtype hepatocellular carcinoma, the dashed lines outline the boundary between normal (left) and tumour (right) regions. The images before and after surface extraction is separated by the blue solid line in HiLo, and the arrows represent the missing parts due to the specimen’s irregular surface. Inset at the bottom right of HiLo shows the photograph of the specimen. **b,c,** Zoomed-in HiLo (left), surface topology (middle), and H&E-stained images (right) of the corresponding dashed regions in a. **d–f,** Zoomed-in HiLo and H&E-stained images of the corresponding dashed regions in a. Scale bar, 100 μm (b–f).

The lung specimen from the first patient (Fig. 5a) is a pathologically confirmed case of large cell carcinoma (LCC). The size of the specimen is 11 mm (length) × 6 mm (width) × 1 mm (thickness), and the total imaging time is within 10 s. Both HiLo image (top of Fig. 5a) and H&E-stained image (bottom of Fig. 5a) outline a clear boundary between normal (right) and tumor (left) regions. The zoomed-in HiLo images (Fig. 5b,c,e,f) show the large tumor cells with vesicular nuclei growing with abundant lymphocytic infiltration, which is the feature of LCC. Leucocytes close to the tumor boundary (Fig. 5d,g) are well identified. Diagnostic features can be quantitatively extracted from the HiLo images. The areas and center positions of 50 lymphocytes and 50 cancer cells are obtained from Fig. 5b. With the localized center position of each cell nucleus, the intercellular distance is calculated to be the shortest adjacent distance to a neighboring cell nucleus. The distributions of nuclear cross-sectional area (Fig. 5h) and intercellular distance (Fig. 5i) are demonstrated, indicating that the cancer cells and lymphocytes can be well distinguished on the basis of these cellular features. In contrast, the specimens from the second and third patients are confirmed cases of lung adenocarcinoma, which presents a different growth pattern compared with LCC. Specifically, the specimen from the second patient is categorized as papillary-predominant adenocarcinoma (Fig. 5j), which is characterized by finger-like papillary architecture with tumor cells lining the surface of branching fibrovascular cores (Fig. 5k,n). The vessels (Fig. 5l,o) and densely packed tumor cells in micro-papillary clusters (Fig. 5m,p) are clearly revealed with remarkably recognizable features comparable to the H&E-stained histological images. The specimen from the third patient is diagnosed with acinar-predominant adenocarcinoma (Fig. 5r), in which tumor cells are arranged in acinar pattern spreading on a fibrotic stroma. The specimen has a size of 11 mm (length) × 5 mm (width) × 5 mm (thickness), with a surface irregularity of ~120 μm. To extract the intact surface profile of the specimen (middle of Fig. 5r), the sample is axially scanned with 10-μm interval, and the total imaging time is within 80 s. The glandular structures are well recognized in Fig. 5s-x, which are well distinguished from papillary adenocarcinoma in terms of the growth pattern.

Finally, a liver specimen is imaged to show the wide applicability of *ex-vivo* fluorescence histology by HiLo microscopy (Fig. 6). The specimen is a pathologically confirmed case of hepatocellular carcinoma (HCC) with scirrhous subtype, in which tumor cell nets are separated by rich fibrous connective tissues. The specimen has a size of 12 mm (length) × 4 mm (width) × 5 mm (thickness), with a surface irregularity of ~200 μm. Similarly, variable focusing is implemented to obtain the specimen’s surface topology (middle of Fig. 6a). The specimen is axially scanned with 10-μm interval, and the imaging is completed within 2 minutes. In view of the specimen’s irregular surface, several FFPE thin slices are sectioned from different tissue depths to obtain the corresponding features between HiLo and H&E-stained images (only one slice shown here, bottom of Fig. 6a). The boundary between normal (left) and tumor (right) regions is readily outlined by HiLo. Zoomed-in HiLo images with their corresponding histological images are demonstrated in Fig. 6b–f. The normal hepatocytes (Fig. 6b) and tumor cells (Fig. 6c) can be well distinguished on the basis of different cellular morphology, although it is not intuitive to diagnose this HCC subtype which featured by intratumoral collagen with only nuclear contrast. With 1.3-μm lateral resolution, small lymphocytes accumulated at the circumference of hepatic vasculature are resolved individually (Fig. 6d,e). These results validates that HiLo enables to evaluate large specimens within point-of-care time frames, demonstrating its great potential as an intraoperative SMA tool that can be used by surgeons and pathologists to provide optimal adjuvant treatment.

## Discussions

HiLo is a promising and transformative imaging technology that enables rapid and non-destructive imaging of large clinical specimens, demonstrating a great potential as an assistive imaging platform that can guide surgeons intraoperatively. To the best of our knowledge, this work demonstrates the first application of using speckle illumination microscopy (i.e., HiLo) for rapid diagnosis of different subtypes of human lung adenocarcinoma and hepatocellular carcinoma, producing images with remarkably recognizable cellular features comparable to the gold standard FFPE histology (Figs. 5 and 6). Although formalin-fixed tumor specimens were imaged in this study to maintain the tissue integrity during transportation from operating room to our lab, fresh tissues are also expected to achieve the similar results (e.g., Fig. 4) by *ex vivo* fluorescence microscopy^33^. Since eosin is weakly bound to cytoplasmic proteins in fresh and hydrated tissues, only nuclear fluorescence channel is implemented in this study to ensure a high image contrast. An optimized two-color fluorescent analog of H&E-staining is reported recently^20^, which can be adapted to this work to facilitate robust intraoperative pathology.

The superiority of HiLo over the gold standard histology is that large surgical specimens can be sufficiently sampled with HiLo microscopy whereas only spot-checked with conventional histology. This is crucial for the improvement of diagnostic accuracy. HiLo should be able to operate at half of the camera framerate (100 fps in our settings) theoretically. The acquisition speed of the current system is mainly limited by the exposure time for each raw fluorescence image, which is ~100 ms with an illumination power of 4 mw, corresponding to an imaging speed of 5 cm^2^/min per channel. Increasing excitation power can further accelerate the acquisition speed by an order of magnitude. For patients who undergo hepatectomy for HCC, the surface area of the specimens can easily reach dozens of square centimeters, which could be imaged within several minutes with the increased imaging speed. The main cause of deviations presented in HiLo and H&E-stained images (Fig. 5a,j,r) is that it’s difficult to orient the specimen in paraffin with an angle exactly the same with that for imaging, which may result in a significantly different area displayed in the H&E-stained slides. However, the histological features which are diagnostically important are still remarkably similar.

As shown in Fig. 2, thick specimens will hamper the optical sectioning strength of HiLo by reducing the in-focus speckle contrast. This can be alleviated by using coarser speckle patterns, which however, will lead to a compromised axial resolution, which is measured to be 5.8 μm with current settings. This axial resolution enables to produce an optical section in lieu of a physical section in conventional slide-based histology. For the lateral resolution, which is currently limited by the sampling pixel size on the detector, will be ultimately determined by the NA of the employed objective lens. Although a lateral resolution of 1.3 μm in this study is capable of resolving densely packed tumor cells as in lung adenocarcinoma (Fig. 5), it is not sufficient to reveal nucleolar structures which are often associated with the degrees of malignancy. Super resolution techniques via synthetic aperture with speckle illumination microscopy^34–36^ could potentially provide a solution to bypass this resolution limit to boost the accuracy of specimen analysis. Another limitation of HiLo is the imaging depth. Similar to SIM, the image SNR will be dramatically decreased when imaging deep into the tissue since the shot noise from background is recorded along with the in-focus signal^37^. This will limit the effectiveness of HiLo for volumetric imaging applications compared with beam-scanning approaches.

The practicality of the HiLo system can be further boosted for clinical translations. First, the narrow DOF (<10 μm with 0.3-NA objective) is not able to accommodate various surface irregularities presented in large specimens, causing the surgical margins to come in and out of focus during the HiLo imaging. To obtain the intact surface profile, variable focusing is required (Figs. 5r, 6a), which will sacrifice the achievable imaging throughput. Recently proposed methods for extended DOF via dynamic remote focusing^38^ or deep learning^39^ could provide a solution to this technical problem, holding a great promise to speed up the whole imaging process by obviating the need for axial scanning. Second, the current system can be developed as an endoscope through a flexible imaging fiber bundle^40–42^. This will enable *in vivo* assessment of remaining tissue after tumor excision, facilitating image-guided surgery which can help surgeons to perform safer and less invasive procedures.

In summary, HiLo is a promising and transformative imaging technology which enables rapid and non-destructive imaging of large clinical specimens, demonstrating a great potential as an intraoperative SMA tool that can be used by surgeons and pathologists to detect residual tumor at surgical margins. It is experimentally demonstrated that HiLo enables rapid diagnosis of different subtypes of human lung adenocarcinoma and hepatocellular carcinoma, producing images with remarkably recognizable cellular features comparable to the gold standard FFPE histology. As a proof-of-concept, this study is limited by the small number of specimens. Future studies will involve more specimens to quantify the diagnostic metrics (i.e., sensitivity and specificity). Moreover, computer-aided diagnosis could be incorporated with HiLo to further improve the efficiency of the current pathological workflow.

## Methods

### Theoretical background of HiLo microscopy

Optical sectioning through HiLo microscopy has been reported in detail^25,26^. HiLo requires two images to obtain one optically sectioned image. A uniform-illumination image (*I_u_*) is used to acquire high-frequency (Hi) components whereas a speckle-illumination image (*I_s_*) is used to obtain low-frequency (Lo) components of the final image. The fusion of these two images will produce a full-resolution optically-sectioned image *I_HiLo_*, which can be calculated as

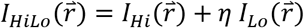

where 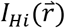 and 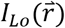 are the intensity distributions of the high- and low-frequency images, and 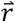 is the spatial coordinates. *η* is a scaling factor that ensures a seamless transition from low to high spatial frequencies, which can be determined experimentally.

It is well known that the intensity of higher frequency components attenuates much rapidly than lower frequency components with the increase of defocus. As a result, high-frequency components are imaged with high contrast only at the focal plane. Therefore, high-frequency components are naturally optically sectioned and they can be extracted from *I_u_* via a simple high-pass filtering

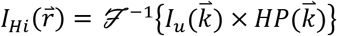

where 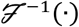 stands for the inverse Fourier transform, and 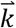 denotes the coordinates in the Fourier domain. *HP* is a Gaussian high-pass filter with a cutoff frequency of *k_c_* in the Fourier domain.

The low-frequency components can be calculated with a complementary low-pass filter *LP* as

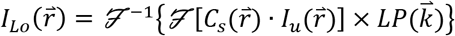

where 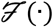 denotes the Fourier transform, and *LP* = 1 – *HP*. Here the speckle contrast 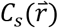 serves as a weighting function that decays with defocus, which enables to distinguish between in-focus from out-of-focus contributions in uniform-illumination images. To eliminate the variations induced by the object itself, the speckle contrast should be evaluated locally on the difference image, which is given by 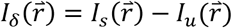 Correct evaluation of the local speckle contrast is crucial to HiLo, and it can be calculated as

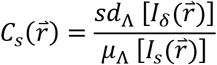

where *sd*_Λ_(·) and *μ*_Λ_(·) represent the standard deviation and mean value calculated over a sliding window with a side length of Λ, which can be determined by Λ= 1/2*k_c_*^28^.

The decay efficiency of *C_s_* can be accelerated by applying an additional band-pass filter to the difference image prior to contrast evaluation. This band-pass filter can be generated by subtracting two Gaussian low-pass filters, which is calculated as

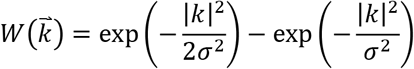

By setting *k_c_* to approximately 0.18σ^26^, the axial resolution of HiLo can be controlled by only changing the parameter σ (e.g., Fig. 3k). The numerical processing can be done in nearly real-time with a workstation with a Core i9-10980XE CPU @ 4.8GHz and 8 × 32GB RAM, and 4 NVIDIA GEFORCE RTX 3090 GPUs.

### Experimental setup

As shown in Fig.1, two laser diodes (CPS 532 and CPS635S, Thorlabs Inc.) emit at 532 nm and 635 nm are implemented in the Epi-mode HiLo system for simultaneous excitation of cytoplasmic and nuclear channel. These two coherent sources are combined by a short-pass dichroic beamsplitter (FF556-SDi01, Semrock Inc.) and propagated through a variable beam expander (BE052-A, 0.5× ~ 2× zoom, Thorlabs Inc.), producing a beam diameter ranging from 1.8 mm to 6 mm (for 532-nm channel). The resulting beam is illuminated on a diffuser (DG10-1500, Thorlabs Inc.) with a beam diameter of ~3 mm, generating a coarse speckle pattern with an averaged grain size of ~4 μm to maintain high image contrast in thick scattering tissues. The diffuser is inserted in a home-built motor-driven rotation mount which can be rapidly rotated at a speed of 8000 rpm to generate the uniform illumination. The illumination patterns are reflected by a dual-edge dichroic beamsplitter (FF560/659-Di01, Semrock Inc.) and introduced to the objective’s back focal plane through a condenser lens (LB1723, *f* = 60 mm, Thorlabs Inc.), ensuring that the speckle size remains relatively constant along the propagation of light. The specimen is illuminated from the bottom with an excitation power of 4 mw, and the excited fluorescence is detected by an inverted microscope which consists of a plan achromat objective lens (Plan Fluor, 10×/0.3 NA, Olympus) and an infinity-corrected tube lens (TTL180-A, Thorlabs Inc.). The fluorescence from two channels are successively filtered by two band-pass filters (FF01-579/34-25 and BLP01-647R-25, Semrock Inc.), and imaged by a monochrome scientific complementary metal-oxide-semiconductor (sCMOS) camera (PCO edge 4.2, 2048 × 2048 pixels, 6.5-μm pixel pitch, PCO. Inc.) which can theoretically reach an imaging throughput of ~800 megabytes/s. In our experiments, the specimen is raster-scanned with 10% overlapping by a 2-axis motorized stage (L-509.20SD00, maximum velocity 20 mm/s, PI miCos GmbH) in two scanning cycles, i.e., one with static diffuser for speckle illumination and the other with rapidly rotated diffuser for uniform illumination. The captured image mosaics are stitched by our developed image stitching software, and the whole system is synchronized by the lab-designed triggering circuits and LabVIEW software (National Instruments Corp.). The exposure time for each raw fluorescence image is set to fill the full dynamic range of 16-bit camera without intensity saturation, which is sample-dependent and generally less than 100 ms in our experiments. The stage settling time between two successive images is set to 10 ms.

### Collection of experimental tissues

The organs were extracted from C57BL/6 mice. The heart, brain, kidney and liver were harvested immediately after the mice was sacrificed. All experiments were carried out in conformity with a laboratory animal protocol approved by the Health, Safety and Environment Office (HSEO) of Hong Kong University of Science and Technology (HKUST).

### Tissue processing

For imaging experiments, the specimens were stained with a cocktail containing 10 μM of TO-PRO3 (T3605, Thermo Scientific In.) and 2% v/v solution of Eosin Y (E4009, Sigma-Aldrich) in 1× PBS for 1 min, and then rinsed in PBS for 30 s, blotted dry with laboratory tissues, and immediately imaged by HiLo. After imaging, the specimens were processed following the standard protocol to obtain the H&E-stained histological images. Specifically, the specimens were processed for dehydration, clearing, and infiltration by a tissue processor (Revos, ThermoFisher Scientific Inc.) for 12 hours, and then paraffin-embedded as block tissues which were subsequently sectioned with 7-μm thickness by a microtome (RM2235, Leica Microsystems Inc.). After that, the thin tissue slices were stained by H&E, and imaged by a digital slide scanner (NanoZoomer-SQ, Hamamatsu Photonics K.K.) to generate the histological images.

## Data availability

All data involved in this work, including raw/processed images provided in the manuscript and Supplementary Information, are available from the corresponding author upon request.

## Code availability

The customized code for HiLo image reconstruction is available at https://github.com/TABLAB-HKUST/.

## Acknowledgments

The Translational and Advanced Bioimaging Laboratory (TAB-Lab) at HKUST acknowledges the support of the Hong Kong Innovation and Technology Commission (ITS/036/19); Research Grants Council of the Hong Kong Special Administrative Region (26203619); The Hong Kong University of Science and Technology startup grant (R9421).

## Author contributions

Y.Z. and T.T.W.W. conceived of the study. Y.Z. and L.K. built the imaging system. Y.Z., and C.T.K.L prepared the specimens involved in this study. Y.Z. performed imaging experiments. C.T.K.L performed histological staining. Y.Z. processed and analyzed the data. Y.Z. and T.T.W.W. wrote the manuscript. T.T.W.W. supervised the whole study.

